# Pelvic irradiation induces behavioral and neuronal damage through gut dysbiosis in a rat model

**DOI:** 10.1101/2023.05.21.541625

**Authors:** B S Venkidesh, Rekha Koravadi Narasimhamurthy, Apoorva Jnana, Dinesh Reghunathan, Krishna Sharan, Srinidhi Gururajarao Chandraguthi, Thokur Sreepathy Murali, Kamalesh Dattaram Mumbrekar

## Abstract

**Background:** Pelvic radiotherapy is the endorsed course of treatment for pelvic malignancies, which frequently cover pelvic primary tumor lesions as well as non-cancerous lymphatic drainage sites in the pelvic area. As a result, pelvic irradiation may indiscriminately cause harm to healthy tissues and organs in the pelvic area in individuals undergoing treatment. Some studies suggest that gut microbial dysbiosis can be correlated with the incidence of radiation-induced toxicities in cancer patients. Since, the consequences were earlier thought to be solely due to the targeted or non-targeted effect of radiation, the role of gut microbiota in the non-targeted effects of radiation and the mechanistic role of the gut-brain axis in the pelvic irradiation scenario is not well explored. Hence, the current study was carried out to explore implication of gut dysbiosis in behavioral and neuronal changes induced by pelvic irradiation.

**Materials and Methods:** 3-4-month-old Sprague Dawley rats were given a single dose of 6 Gy pelvic irradiation. Fecal samples of control and treated mice were collected at different timepoints to assess microbial abundance and diversity using 16S rRNA-based metagenomic sequencing. Behavioral analysis, histological analysis of intestine, brain and gene expression analysis of brain hippocampus was performed to ascertain the indirect impact of microbial dysbiosis on cognition.

**Results:** Following pelvic irradiation, significant microbial dysbiosis and behavioral alterations were observed with distinct changes in the microbial diversity and a significant decline in the locomotor effect and anxiety level at each time point following radiation. Histological analysis revealed a significant reduction in villus distortion as well as a significant decrease in neuronal cells, matured neurons, and an increase in reactive astrocytes, suggesting that pelvic irradiation promotes neuroinflammation. Gene expression analysis revealed a significant reduction in neural plasticity. Altogether, this study demonstrated that gut dysbiosis caused by pelvic irradiation alters behavior, intestinal morphology, integrity, and brain neuronal maturation, as well as lowers the levels of neural plasticity expression.

**Conclusion:** Current study provides evidence for the influence of gut dysbiosis on pelvic irradiation induced cognitive impairment in a rat model.

## INTRODUCTION

Over 50% of cancer patients undergo radiotherapy at some point during their treatment, and it is a pillar of the modern management of malignancies (Baskar et al., 2012). Up to 75% of cancer patients undergoing radiotherapy experience radiation-induced adverse effects (Zimmerer et al., 2008). The prevention of side effects linked with radiotherapy has received much attention due to the associated improvement of survival rate among the patients (Barrazuol et al., 2020). Additionally, radiotherapy for abdominal and pelvic malignant tumors causes symptoms of acute intestinal toxicity in 60–90% of patients, which significantly lowers their quality of life and disrupts their treatment regimen (Kumagai et al., 2018). Ionizing radiation (IR)-mediated gastrointestinal (GI) toxicity is primarily defined as clonogenic cell death and apoptosis in the crypt cells which results in the inadequate repair of villus epithelial cells, destruction of the mucosal barrier resulting in mucositis, and a notable reduction in the compensatory inflammatory responses (Wang et al., 2004). These radiation-induced changes in the GI microenvironment results in the alteration of gut microbiota (Wang et al., 2021).

The human gastrointestinal tract (GIT) is colonized by diverse gut microbiota that play a major role in host immunity, defense against infections, as well as a wide variety of other physiological tasks that are beneficial to the host (Thursby et al., 2017). The diversity and abundance of gut microbiome can be influenced by a variety of external factors including their lifestyle patterns, radiation exposure, food habits, nutritional status, and medical history (Joseph et al., 2020). Various clinical studies have reported alterations in gut microbiome in association with a wide variety of cardiometabolic and chronic diseases, such as obesity, type 2 diabetes, atopic diseases, cardiovascular diseases, hypertension, anxiety, depression, bowel diseases, diarrhea, constipation, and brain disorders such as Parkinson’s and Alzheimer’s disease (Blutt et al., 2018). The finding that functional and inflammatory gastrointestinal illnesses can contribute to psychiatric comorbidity, including depression and anxiety in up to 80% of patients, lends support to the hypothesis that changes to the microbiota can affect central nervous system (CNS) functions (Keightley et al., 2015). Microbes communicate with brain, via various mechanisms involving vagus nerve, short chain fatty acids (SCFAs), cytokines, hormones, γ-aminobutyric acid (GABA), acetylcholine and several other neurotransmitters (Cryan et al., 2019). This bidirectional communication network that connects the host gut and brain activities is known as microbiota-gut-brain axis (Raskow et al., 2016). Cui et al. (2017a) using a mice model showed that the total abdominal irradiation causes a characteristic shift in gut microbiota with corresponding behavioral changes providing direct evidence of the pathogenic mechanisms behind the distant cognitive impairment brought on by abdominal and pelvic radiation. Remarkably, Luo et al. (2020) found that in mice administered with whole brain irradiation (following depletion of native microbial flora), treatment with probiotics could improve spatial memory function. This highlights the importance of gut flora in reversing the cognitive impairment brought on by radiation-induced brain injury. Further, studies have proven that intestinal flora depletion following radiation therapy may serve as a protective modulator against radiation-induced brain damage (Luo et al., 2022). Also, the gut damage due to radiation contributes significantly to gut dysbiosis due to the associated inflammation (Stringer et al., 2011), loss of intestinal integrity (Ulluwishewa et al., 2011) and increase in abundance of more pathogenic or harmful gut bacteria (Jian et al., 2021).

The current study was designed to understand the effect of pelvic irradiation on neurotoxicity manifested through gut-brain axis and whether this could be correlated with changes in gut microbial composition. Towards this, we have analyzed the changes in gut microbiota at various time points following a single dose of pelvic irradiation and its effect on the brain through behavioral, neuronal, and gene expression studies in a rat model.

## MATERIALS AND METHODS

### Animal study design

All experiments were conducted in compliance with guidelines that were authorized by the Institution of Animal Ethics Committee (**IAEC/KMC/84/2020**). The maintenance of animals was carried out under standards established by the WHO, Switzerland, and the Indian National Science Academy, New Delhi. Male Sprague Dawley rats (3-4 months old) weighing 200-250 g were fed under controlled environmental conditions (23 ± 2 °C), humidity (50 ± 5 %), light (12:12 hr of light/dark cycle), and with continuous access to standard feed and sterile water. Rats were divided into control (no treatment) and radiation groups (n=6/group). Animals of irradiation group were restrained in plexi glass restrainers and received a single dose of 6 Gy of radiation with a dose rate of 306 MU/min to the pelvic region using Elekta Medical Linear Accelerator (Versa H, Stockholm, Sweden) delivering 6MV photon energy. The pelvic irradiation was done at Shirdi Saibaba Cancer Hospital and Research Centre, Kasturba Hospital. Fecal samples (n=3/timeline/group) were collected on 0, 7 and 12 days post-irradiation, followed by the behavioral experiments which were performed between 14^th^ and 16^th^ day (Figure 1).

**Figure 1:**
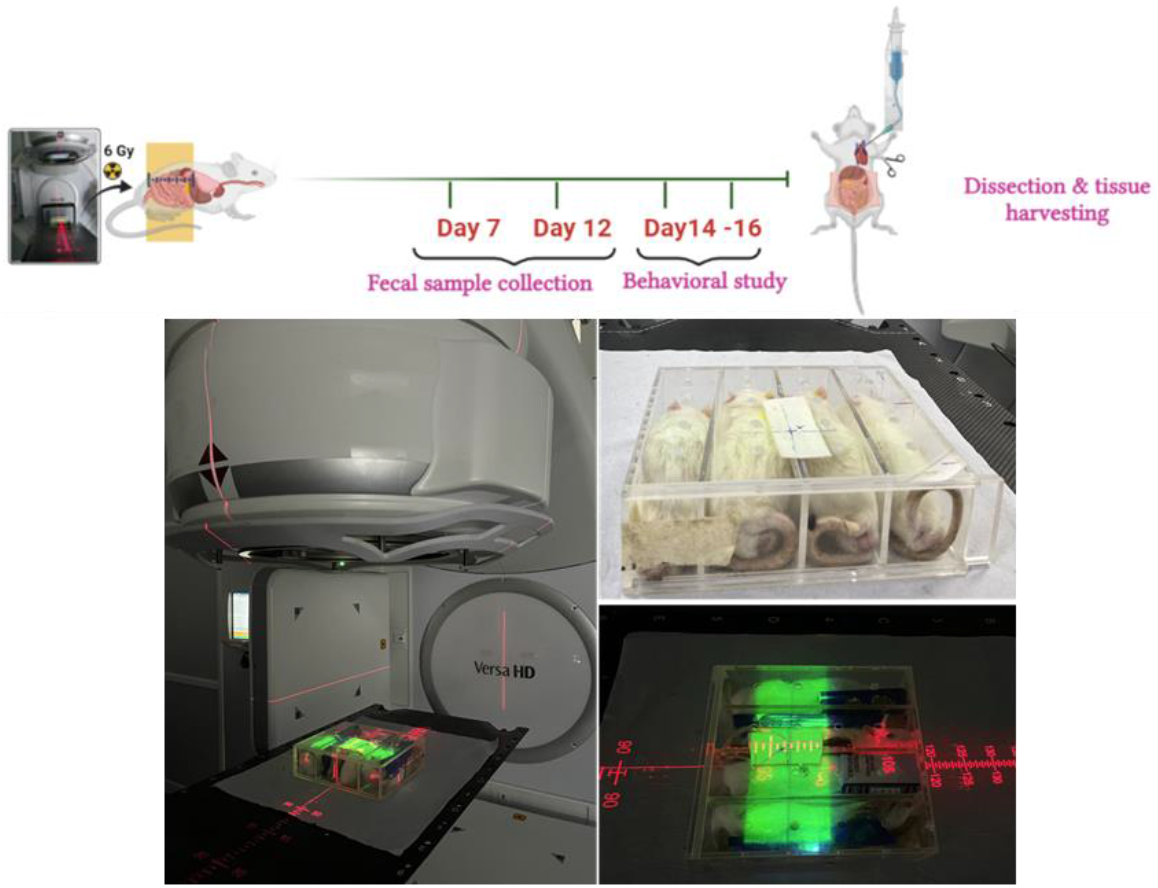
The time line for the study and radiation exposure setup.

### Microbial enumeration

#### Fecal sample collection and DNA isolation

For the collection of fecal samples without external contamination, animals were placed in individual autoclaved cages and allowed to defecate. The fecal samples were then collected in an autoclaved 1.5 mL polypropylene tube and stored in -80°C until further analysis. Fecal DNA was isolated with standard phenol-chloroform methods (Sambrook et al., 2001). Briefly, 200 mg of frozen fecal samples were weighed and 800 μL of lysis buffer (TES: Tris-EDTA-sodium dodecyl sulphate SDS) was added sulphate) buffer, 40 μL of 20% of SDS, and 10 μL of proteinase K (10mg/mL) were added. The samples were homogenously mixed, vortexed and samples were incubated at 70°C for 20 minutes in a water bath. An equal volume of phenol was added, then centrifuged at 9408 ×g (10,000 rpm) for 10 minutes at 4°C. To the aqueous phase, an equal volume of chloroform: isoamyl alcohol (24:1 ratio) was added and then centrifuged as before. The aqueous phase was transferred into a fresh tube to which double the volume of 100% ethanol and 1/10^th^ of 3 M sodium acetate was added, inverted gently, and placed in -80°C for 2 hours. The samples were then thawed at room temperature and centrifuged. RNase treatment was performed by addition of 100 μL of 1 M Tris-EDTA (TE) and 20 μL of RNase (10 mg/mL) to the pellet followed by incubation at 37°C for 1 hour in a water bath. Further, 0.7X isopropanol and 1/10^th^ of 3M sodium acetate was added to the tubes, gently inverted, and then centrifuged. Resulting DNA pellet was washed with 500 μL of chilled 70% ethanol, centrifuged at 9408 ×g for 10 minutes at 4°C and air-dried on the bench top. DNA pellets were dissolved in 40 μL of TE buffer and stored at -20°C until further use. The concentration and the purity of isolated rat fecal DNA was determined using Nanodrop spectrophotometer (Thermo Fisher Scientific, USA).

#### 16S rRNA metagenomic sequencing

The concentration of the isolated fecal DNA samples was checked with a Qubit 2.0 fluorometer (Invitrogen, Life Technologies, USA). The 16S rRNA metagenomic sequencing was performed as described in Jnana et al. (2020). Briefly, 50 ng of DNA sample from each group; control, D7R and D12R (7^th^ day post-irradiation and 12^th^ day post-irradiation, respectively) (n=3/time line/group) was processed for 16S rRNA amplification with two primer sets directed at the V2, V3, V4, V67, V8, and V9 hypervariable regions of 16S rRNA. Using a magnetic bead-based PCR purification (Agencourt AMPure XP), the 16S amplicons were purified and pooled (Beckman Coulter, Life Sciences, US). Ion Plus Fragment Library kit™ and Ion Xpress Barcode Adapters™ were used to end repair, ligate and multiplex using barcodes. Using the DNA high-sensitivity kit in the 2100 Bioanalyzer (Agilent Technologies, USA), the library quality and quantity of the appropriate fragment length (250 bp) were evaluated (Agilent Technologies, USA). Each library was diluted to achieve a DNA concentration of 26 pM. Ion One Touch 2 and Ion One Touch ES systems were used to create equimolar template positive ISP libraries and loaded on to 318v2 chip and sequenced using Ion Personal Genome Machine via 400bp sequencing chemistry (Thermo Fisher Scientific, USA). Following sequencing, Torrent Suite v5.0 (Thermo Fisher Scientific, USA) was used to carry out base calling and demultiplexing of the sequencing runs using the default settings. Further, the OTU (operational taxonomic units) reads were processed to calculate the relative abundance (RA) using the formula: RA = (number of reads for a particular OTU/total number of reads) × 100. Then, the metadata and taxonomy files were manually generated according to the species present in the OTU table. All the files were exported to excel worksheet and converted to .txt and .csv formats for further processing. Microbiome Analyst v2, a web-based tool, was used for downstream analysis of microbiome data (https://www.microbiomeanalyst.ca/MicrobiomeAnalyst/upload/OtuUploadView.xhtml). The marker gene data profiling (MDP) module was selected to analyze the marker gene survey data (Chong et al., 2020). After, the data normalization and filtering, relative abundance, and alpha and beta diversity were calculated and heatmap clustering was performed. Venny 2.0 was used to visualize the overlap between microbial members of each group (control and treatment – D7R, D12R) (https://bioinfogp.cnb.csic.es/tools/venny/). All the analyses were performed at family and genus levels for samples obtained at different timepoints (C, D7R and D12R) in both control and radiation groups.

### Data availability

Metagenomic raw sequencing data of this study is available under the Bioproject identifier PRJNA967983.

### Behavioral Assessment

#### Elevated Plus Maze Test (EPM)

On the 15^th^ day post-irradiation EPM test was performed on rats to analyze anxiety-like behaviour by measuring the time spent on exploring a novel environment and avoid elevated and open spaces. The apparatus consists of two open (stressful) and closed (protective) elevated arms. Investigation of the open arm (OA) was conducted under indirect dim light. Initially, the animal is placed at the intersection, facing OA. After testing the behavior of each rat, the maze was cleaned with alcohol. Parameters measured include the number of entries in the OA and time spent in the OA (Tillmann and Wegener et al., 2019).

#### Novel Object Recognition Test (NOR)

NOR test investigates the learning and memory in rats. NOR mainly detects the tendency to investigate an unknown object when exposed to either a familiar or a novel object. The three days of experimental flow involves habituation, training, and test session, mainly reflecting the function of dorsal hippocampus. Exploratory behavior scoring includes time spent in the center square and the total investigation time at the novel object. Analysis was made using video recordings (Lueptow et al., 2017).

#### Histopathological Analysis

Following the behavioral experiments, animals were deeply anesthetized and perfused with 0.9% saline. After euthanisation, the intestine and brains were removed and fixed in 10% neutral buffered formalin. Following routine processing in paraffin, sections of the intestine, brain hippocampus, and cortex region (n=3/group) were cut at 5μm thickness in a rotary microtome (Leica RM2125RT, Wetzlar, Germany) for histological analysis (Venkidesh et al., 2023). The staining protocol was performed following laboratory standardised method (Venkidesh et al., 2023). Images were captured using a light microscope (OLYMPUS CKX53, Tokyo, Japan) at 40X magnification for visualizing the H&E-stained intestinal sections, 100X magnification for visualizing the PAS-stained goblet cells, and 600X for the immunohistochemistry-stained cells.

#### Hematoxylin and eosin (H&E) staining for intestinal morphology

Intestinal morphology like villous-crypt architecture, height was analyzed using the H&E staining. Intestinal (jejunum) parameters included the villus height (vertical distance from the villous tip to the villous-crypt junction level), villous distortion, and crypt architecture depth. A total of 10 villi and 10 crypts per section were analysed (Messora et al., 2013; Foureaux et al., 2014).

#### Periodic-Acid staining (PAS) for intestinal integrity

PAS staining was performed to assess the goblet cells in the intestine. Slides containing sections were oxidized in 1% periodic acid for 10 mins and then immersed in Schiff’s reagent for 10 minutes and then counterstained with hematoxylin for 1-2 minutes (Li et al., 2019).

#### Nissl staining for neuronal viability

Nissl staining is an effective method to analyse the morphology and pathophysiology of neural tissue. Staining was performed using laboratory standardised method (Venkidesh et al., 2023). Images were captured under 200X magnification using a light microscope (Olympus, IX-HOS, Tokyo, Japan).

#### Immunohistochemistry (IHC) for neuronal maturation and inflammation

This experiment was performed for the purpose of analyzing the neuronal differentiation to assess the functional state of neurons and astrogliosis, which leads to inflammation in tissues (Sofroniew et al., 2015). The protocol was performed following laboratory standardised method (Venkidesh el al., 2023). The sections were immunostained by primary antibodies like neuronal nuclear protein (Neu-N) (Invitrogen, California, USA, 1:500 dilution) (Gusel’Nikova et al., 2015), glial fibrillary acidic protein (GFAP) (Invitrogen, California, USA, 1:2000 dilution) and with HRP (horse radish peroxidase)-conjugated goat anti-rabbit IgG (secondary antibody) (Invitrogen, California, USA, 1:1000) for 1hr, treated with DAB (3,3′-Diaminobenzidine) chromogen (Dako, North America), counterstained with hematoxylin, and mounted using Dibutylphthalate Polystyrene Xylene (DPX). Images were captured using a light microscope (Olympus, IX-HOS, Tokyo, Japan).

#### Hippocampal Gene expression

The part of brain hippocampus tissue (n=3/group) was weighed and homogenized in 500 μL (for 50 mg of tissue) Trizol reagent and stored at -80°C until further use. Total RNA was isolated with Trizol reagent as per the manufacturer’s protocols (Thermo Fisher Scientific, USA). The homogenized tissue was thawed on ice and centrifuged at 13548 ×g (12,000 rpm) for 5 minutes. Pellet was mixed with 200 μL of chloroform and placed on ice for 15 minutes followed by centrifugation at 4°C, 13548 ×g (12,000 rpm) for 15 minutes. The upper aqueous phase was mixed with an equal volume of isopropanol and stored at −20°C for 2 hours. Samples were centrifuged at 4°C, 13548 **×**g (12,000 rpm) for 10 minutes and the supernatant was discarded. Pellets were washed with 1 mL of 70% ethanol and centrifuged at 4°C, 5724 **×**g (8000 rpm) for 5 minutes. RNA precipitating at the bottom of the tube was dried at room temperature for 5-10 minutes and dissolved in 20-30 μL DEPC-treated deionized water. The concentration and purity of RNA was detected using Nanodrop spectrophotometer (Thermo Fisher Scientific, USA), and integrity was assayed with gel electrophoresis. With the aid of a Thermo Scientific RevertAid First Strand cDNA Synthesis Kit, extracted RNA was reverse transcribed into cDNA (Thermo Scientific, USA).

Using the Power SYBR™ Green PCR Master Mix, quantitative real-time PCR (qRT-PCR) was used to analyze the expression of target genes related to neuromodulation and neuroplasticity. The sequences of primers used to analyze the gene expression were as follows: *Bdnf* F (5’-CCCTGGCTGACACTTTTGAG-3’) and R (5’-GAAGTGTACAAGTCCGCGTC-3’), *Nmda2* F (5’-GCACACGAGTCAGAGGTTTC-3’) and R (5’-AGAGTCCCCTTCTGTCTTGC-3’). *Gapdh* was used as the housekeeping gene with the primers: *Gapdh* F (5’-GTATGACTCTACCCACGGCA-3’) and R (5’-CCCCATTTGATGTTAGCGGG-3’). qRT-PCR was performed in Quant Studio 6 Pro (Applied Biosystems, USA). The data obtained was analyzed to determine the relative expression of target genes with *Gapdh* serving as the reference gene.

#### Statistical Analysis

A one-way ANOVA test with multiple comparison analysis was performed using GraphPad Prism version 8.0, and p<0.05 was considered as statistically significant.

## RESULTS

### Microbiome diversity at family and genus level

In this study, we obtained a total of 14,76,608 16S rRNA sequence reads from the rat fecal DNA collected at different time points after radiation treatment. Following quality control, a total of 136,008 sequenced reads were mapped to the control sample, 76,363 sequenced reads were mapped to the D7R sample and 48,463 sequenced reads were mapped to the D12R sample. A total of 71 OTUs at the family level and 46 OTUs at the genera level belonging to 10 different phyla were identified from the fecal samples collected at different time points. The OTUs at genera level were found to be shared among the control, D7R and D12R includes *Roseburia, Blautia, Tyzzerella, Paraprevotella, Faecalibacterium, Parabacteroides, Desulfovibrio, Bifidobacterium, Anaerobiospirillum, Aeromonas, Eubacterium, Parasutterella, Bacteroides, Clostridium, Ruminococcus, Prevotella*, and *Lactobacillus*. We observed that 11.7% and 37% of OTUs were unique to samples collected at D7R and D12R respectively. D12R showed the highest richness at the genera level (Figure 2A).

**Figure 2.**
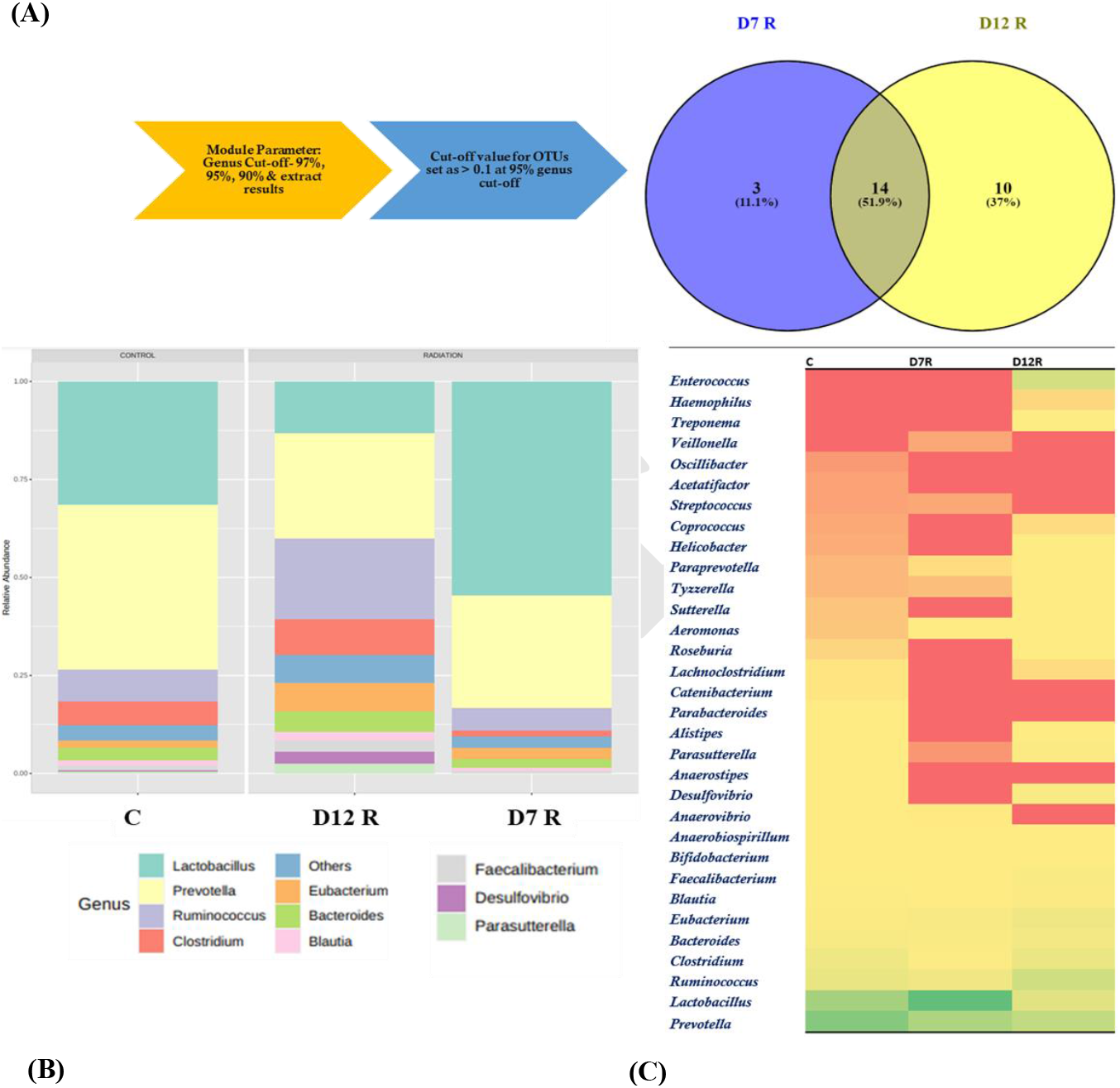
Changes in gut microbiota post pelvic radiation. (A) The module parameters applied throughout the raw data processing & the Venn diagram showing shared and unique OTUs at genus level for D7R and D12R samples (B) Relative abundance at bacterial genus level from different samples and (C) Heat map clustering of microbes at genus level across samples (C – control; D7R – 7 days post-irradiation; D12R – 12 days post-irradiation)

Based on the relative abundance values, the following genera were found to the dominant across both the groups: *Prevotella, Ruminococcus, Lactobacillus, Clostridium, and Bacteriodes* (Figure 2B). However, a characteristic change in the abundance level of these genera was observed in the timeline D12R, indicating highest changes in gut microbiota at 12^th^ day post-irradiation.

### Clustering of rat gut microbiota in the different timelines at family and genus level

Based on heat map clustering analysis, we could discern clear differences in bacterial assemblage (genera level) associated with samples obtained at different time points. The bacterial assemblage isolated from control and D7R samples were clustered together, compared to the assemblage from D12R sample. The major bacterial genera found in D12R that were significantly abundant included *Parabacteriodes, Sutterella, Desulfovibrio, Ruminococcus, Treponema, Alistipes, Parasutterella, Helicobacter, Eubacterium, Tyzzerella*, which belonged to the families Helicobacteraceae, Spirochaetaceae, Sutterellaceae, Desulfovibrionaceae and Tannerellaceae (Figure 2C).

### Intestinal morphology and integrity

Morphological parameters like villous distortion, crypt architecture, and villous height were analyzed and we found that pelvic irradiation significantly damaged the villous architecture (p<0.01) and crypt architecture (p<0.05) compared to the control group. However, no significant difference was observed in radiation-induced villi height (Figure 3A). Further, PAS staining showed that radiation exposure significantly reduced the number of mucin-producing goblet cells (p<0.05) (Figure 3B). Altogether, a single dose of 6 Gy of pelvic irradiation resulted in significant changes in intestinal morphology and integrity.

**Figure 3.**
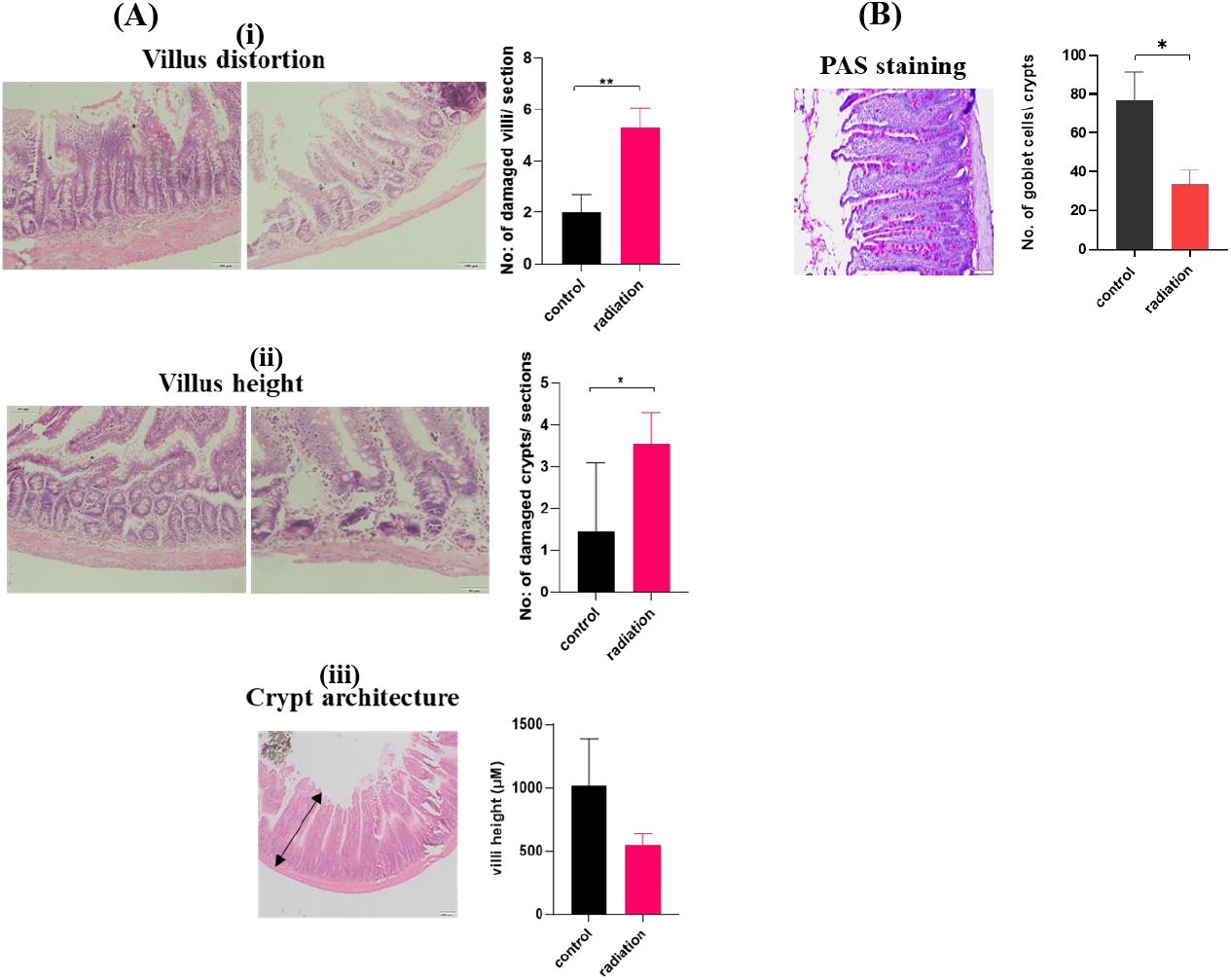
(A) Intestinal morphological analysis using hematoxylin and eosin (H&E) staining. Representative images showing H&E-stained intestinal sections of Control and Radiation groups. (i) villus distortion, (ii) villus height and (iii) crypt architecture; (B) Intestinal integrity analysis after pelvic radiotherapy. Representative image showing PAS-stained intestinal section. Data shown as mean ± SEM (n=3); P values: *<0.05, ** <0.01

### Neuronal survival, neuroinflammation and neuronal maturation

The neuronal survival analysis was performed on Dentate Gyrus (DG) and Cornu Ammonis 2 (CA2) regions of hippocampus, responsible for memory and cognition. Pelvic irradiation increased the number of pyknotic neurons in the DG region (p<0.0001) and CA2 region (p<0.01) (Figure 4A). Neuronal survival analysis indicated that pelvic irradiation reduced neuronal viability. In comparison to the control, a significant increase in reactive astrocytes was seen in the CA1, CA2, cortex, and DG regions of the brain in the radiation group (p <0.05) (Figure 4B), suggesting that pelvic radiation could induce neuroinflammation. We also found that pelvic radiation reduced (p<0.001) the matured neurons in the CA1, CA2 and CA3 region of the rat hippocampus (Figure 4C). This suggests that the low dose radiation in the pelvic region altered the neuronal survival, maturation, and induced neuro-inflammation.

**Figure 4.**
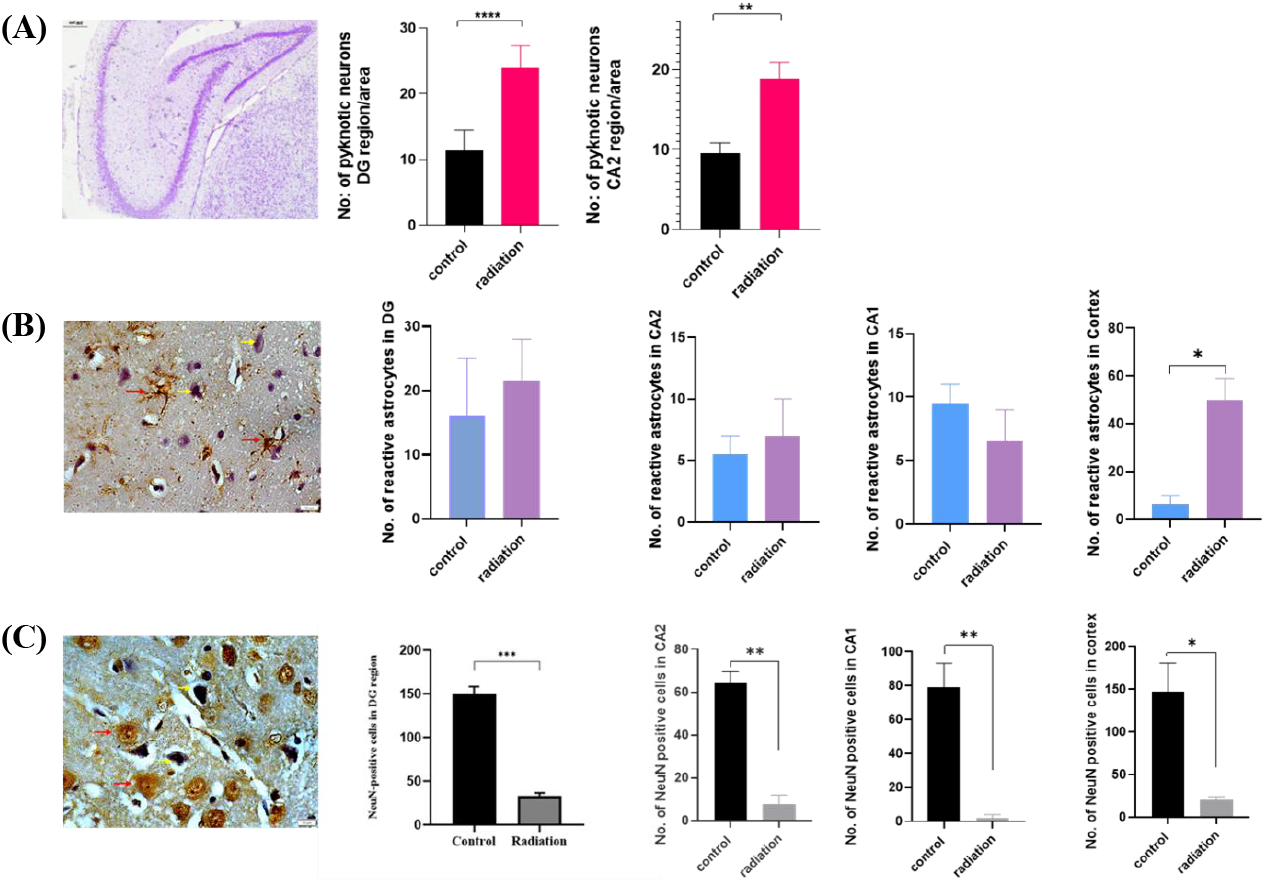
Effect of pelvic irradiation on neuronal survival, neuroinflammation and neuronal maturation (A) Representative image showing Nissl-stained hippocampus (DG and CA2) regions and the number of pyknotic cells in DG and CA2 regions. (B) Representative image showing the reactive astrocyte (GFAP stained with DAB chromogen) and reactive astrocytes in DG, CA2, CA1 and Cortex regions. (C) Representative image showing the Neu-N-stained matured neuron and the number of matured neurons in DG, CA2, CA1 & Cortex region. Data shown as mean ± SEM (n=3); P value *<0.05, ** <0.01, *** <0.001, **** <0.0001 (Red arrow shows the positively stained cells and the yellow arrow shows hematoxylin-stained cells).

### Memory and anxiety

In our study, a single dose of 6 Gy of pelvic irradiation resulted in a reduction in the exploratory behavior of animals in the irradiated rats (p<0.05) (Figure 5). However, animals exposed to radiation did not show any changes in the memory, based on the time spent on the novel object analysis test (Figure 5A). Similarly, we did not observe any alterations in anxiety levels as measured by the number of entries made by rats in the OA and time spent in the OA (p>0.05) (Figure 5B).

**Figure 5.**
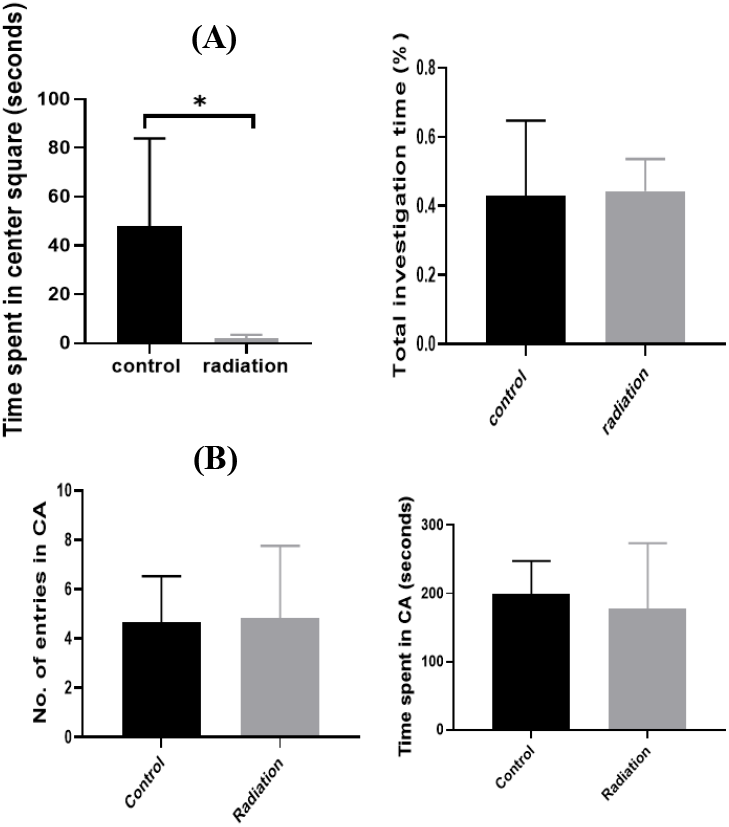
Effect of pelvic irradiation on memory and anxiety (A) Novel Object Recognition (NOR) test results after radiation exposure, (B) Elevated Plus Maze test results after radiation exposure; Data indicated as mean ± SEM (n=6), p value* <0.05.

### Gene expression

To further elaborate on the indirect effect of pelvic irradiation on neuronal plasticity and neuromodulation, the expression of *BNDF* and *NMDA2* genes in the brain tissue was analyzed. We found the gene expression level of *NMDA2* was reduced in the radiation group (p<0.05) (Figure 6). However, the expression level of *BDNF* was not affected by pelvic irradiation (p>0.05). This suggests that a single dose of pelvic irradiation could significantly reduce the expression level of neuronal plasticity but had no effect on the gene responsible for neuromodulation.

**Figure 6.**
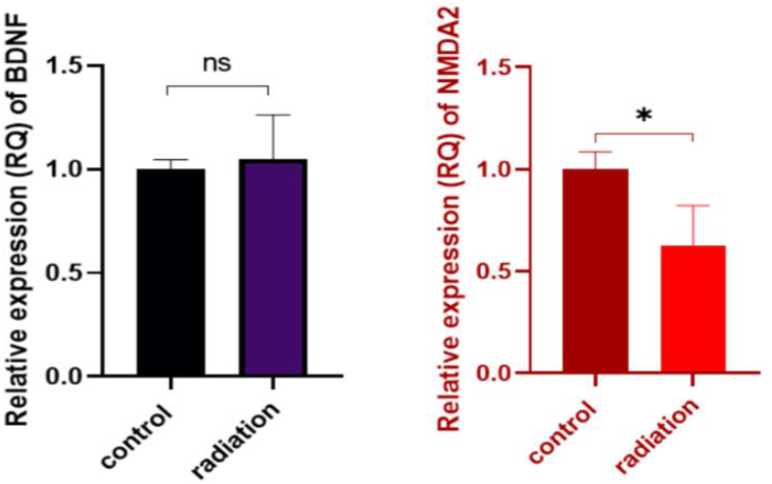
qRT-PCR analysis of BDNF and NMDA2 in brain hippocampus tissue collected from control and pelvic irradiated rats. Data were analysed using one way ANOVA, n=3, p value* <0.05.

## DISCUSSION AND CONCLUSION

Pelvic irradiation-induced neurotoxicity through gut microbiota axis is poorly understood, as it can have significant impact on the quality of life of patients undergoing radiotherapy. Reports indicate that radiation-induced damage in cancer patients could be associated with gut microbial dysbiosis and radiation is one of the causes of gut microbial dysbiosis (Wang et al., 2019, Joseph et al., 2020). The gut dysbiosis not only impacts the tumor response to radiotherapy but also has a significant influence on the normal tissue response (Liu et al., 2021). In the present study, pelvic radiation of 6 Gy resulted in alteration of the gut bacterial composition, as observed by 16S rRNA sequencing of the gut microbiome. In an earlier study, Cui et al. (2017) showed a direct association between the microbiome and radiation sensitivity in radiation enteritis patients and observed that 6.5 Gy of whole-body radiation altered the community composition of the intestinal microbiota in C57BL/6 mice. Similar findings were made by Zhao et al. (2020), who demonstrated that abdominal radiation dramatically decreased the diversity of the gut microbiota in mice and caused a shift in the balance of their intestinal flora. Additionally, it was observed that the relative abundance of bacteria belonging to the genus *Lactobacillus* was significantly decreased after radiation exposure, indicating that they are some of the most radiosensitive bacteria. In another study, in mice exposed to 5 Gy of total body radiation (TBI), the fecal microbial diversity decreased on day 3 and was found to return to original levels on day 30 post-irradiation (Goudarzi et al., 2016). Li et al. (2019) and Lu et al. (2019) studied the composition of the gut microbiota after exposure to radiation. The microbiome was significantly altered after 3.5 days post 9 Gy TBI (Li et al., 2019) while a similar observation was made by Lu et al. (2019) when they studied the microbiome composition 5 days after 15 Gy whole abdominal irradiation. In both the cases, there was a significant decrease in the relative abundance of Bacteroidetes and an increase in the abundance of Proteobacteria; however, the alpha diversity of the gut microbiota was unaffected (Li et al., 2019; Lu et al., 2019).

We found that samples collected on day 12 post treatment with 6 Gy of pelvic irradiation (D12R) was significantly enriched in abundance of certain bacterial genera which included *Parabacteroides, Sutterella, Desulfovibrio, Ruminococcus, Treponema, Alistipes, Parasutterella, Helicobacter, Eubacterium*, and *Tyzzerella*. Earlier studies have identified a link between *Parabacteroides* and metabolic syndrome, inflammatory bowel disease, and obesity (Cui et al., 2022). Further, the bacterial genera *Parabacteroides* and *Bacteroides* are closely associated with gut inflammation (Lopetuso et al., 2018). Similarly, *Sutterella* has been reported to be found in higher abundance in radiation-induced animal models and is reported to play a role in inflammation damage (Gerrassy et al., 2018). Furthermore, *Sutterella* spp. has been associated with Autism, Down syndrome, and metabolic syndrome in several earlier studies (Williams et al., 2012; Wang et al., 2013; Biagi et al., 2014; Lim et al., 2016). *Treponema* is also found to be significantly enriched upon radiation treatment and is considered a biomarker of radiation induced damage.

The gastrointestinal tract is a very complicated organ that carries out a variety of dynamic physiological and biological processes. The host’s development, health, and disease are all influenced by the gut microbiota in addition to positive host communication. It can support mucosal immune homeostasis, intestinal epithelial regeneration, and barrier integrity (Parker et al., 2018). According to Shadad et al. (2013), intestinal damage from radiation exposure occurs either directly through intestinal stem cell apoptosis or indirectly through endothelial cell-mediated intestinal stem cell malfunction and apoptosis (Paris et al., 2001). Additionally, prior research has demonstrated that the digestive tract is highly radiosensitive and that exposure to sublethal levels of gamma radiation can cause intestinal damage, including the loss of crypt depth, villus height, and several crypts, as well as signficant decrease in intestinal epithelial cell migration (Kumar et al., 2018; Venkateswaran et al., 2019). In our study, we found that the pelvic radiation could result in significant distortion in villus structure, loss of crypt architecture, and villus height.

The hippocampus region of the brain plays a major role in memory and learning (Sekeres et al., 2018) and hence we evaluated the indirect effect of radiation on the hippocampal region of brain at structural as well as functional level. The well studied biological mechanism for radiation-mediated damage is through DNA damage, which disrupts numerous signal pathways involved in the cell cycle, apoptosis, and stress response (Watters et al., 1999). Ionizing radiation can be especially harmful to the hippocampus. According to previous reports, learning and memory problems brought on by radiation are accompanied by an increase in hippocampus apoptosis and a decrease in neurogenesis (Karimi et al., 2018). It is conceivable that cellular elements implicated in both hippocampus apoptosis and neurogenesis may be the cause of hippocampal vulnerability (Tada et al., 2000). In our study, we observed that pelvic radiation can significantly increase the pyknotic (dead) neurons in DG and CA2 region of the hippocampus, which supports our hypothesis that radiation has an indirect effect on memory and cognition. Further, the study focused on the astrocyte cytoskeletal protein GFAP which is a distinctive marker for astrocytes. The level of GFAP expression can be utilized to study the severity of astrocyte injury because activated astrocytes can significantly raise their levels (Zhou et al., 2020). The increased levels of activated astrocytes in the hippocampus region indicated that that the pelvic irradiation could significantly contribute to increased levels of reactive astrocytes in the CA1, CA2 and DG regions of the hippocampus which could further lead to astrocytosis and neuroinflammation. In our study, we observed that 6 Gy of irradiation resulted in an enlarged cell body, increased cell branching, and thickened protrusions. However, preclinical studies have shown that after a single dosage of 20-45 Gy of radiation, GFAP levels can increase over a period of 120 to 180 days, but not when exposed to lower doses (2 or 8 Gy) (Chiang et al., 1993).

Radiation-induced brain injury may result in molecular, cellular, and functional changes to the brain’s neuronal, glial, and vascular compartments. However, the indirect effect of radiation on the brain have been not yet well understood. The hippocampus has been the subject of the majority of studies because of its crucial role in memory and adult neurogenesis. Interestingly, we observed that pelvic radiation could significantly reduce the matured neurons in the hippocampus; however, the role of gut dysbiosis in reduced neuronal maturation, impaired memory, and neurotoxicity, needs further validation. However, gut dysbiosis studies that resulted in an increase in certain bacterial genera like *Parabacteroides, Bacteroides, Clostridium, Sutterlla, Treponema*, etc. have shown that this can significantly contribute to impaired brain function, as was observed in our study.

Cryan et al. (2019) suggested that the gut microbiota could regulate the functioning of CNS, by changing the behavior, memory, and cognition, whereas the brain can activate signaling pathways that affect immune and metabolic function as well as host behavior. Interestingly, several earlier studies have shown that gut dysbiosis could potentially lead to reduced exploratory behavior (Berick et al., 2011), memory dysfunction (Gareau et al., 2011), anxiety-related behavior, impaired social cognition (Hoban et al., 2016) and decreased locomotory activity (Ceylani et al., 2018). Similar to these reports, we also observed in the current study that radiation-induced bacterial dysbiosis to cause a significant reduction in exploratory behavior in rats.

It is well known that radiation-induced changes in neuronal changes as well as impaired expression of genes are involved in memory and cognition. Also, gut microbiota plays a vital role in maintaining brain function. However, impairment in such factors can be associated with gut dysbiosis. The expression of 5-hydroxytryptamine (5-HT) receptors, neurotrophic factors (such as BDNF), and N-methyl-d-aspartic acid (NMDA) receptor subunits in the hippocampus, as well as myelination in the prefrontal cortex, are also altered by gut microbial dysbiosis, which impairs social cognition (Bercik et al., 2011; Hoban et al., 2016). In our study, we observed that pelvic radiation-induced gut dysbiosis could lead to a decrease in BDNF level and a significant reduction in the expression of NMDA2.

In summary, our work reveals that a single dose of 6 Gy of pelvic radiation administered in a rat model showed a characteristic and significant shift in gut microbiota and induced significant damage in intestinal tissue. Interestingly, our data support our hypothesis that the non-targeted effect of pelvic radiation could include a significant loss of neuronal survival, maturation, and exploratory behaviour in rats. Further, it contributed to reduced levels of expression of genes that regulate neuronal plasticity. In summary, our current study indicates that pelvic radiation can induce significant changes in the brain through the gut microbiome-brain axis. However, one of the limitations of the study is that the animal model used could not explain the neuro-metabolomic pathway and underlying mechanisms that contribute to neuronal damage, which we plan to investigate in the future.

## Acknowledgments

This study was supported by MAHE intra-mural grant (2019). VBS would like to thank MAHE for the Dr. TMA Pai Ph.D. Fellowship. The authors would like to express their gratitude to Mrs. Sandhya for assistance with 16S sample processing, Ms. Rekha K N for helping in behavioral experiments and histopathological experiments, Mrs. Kirthi Kulal and Mrs. Anjali Warrier for their contributions to the molecular biology section, Mr. Jackson Rodrigues for helping and troubleshooting in animal experiments, Mr. Ankit Tanwar for giving suggestions during the 16S data processing. We express our sincere gratitude to the technical support team for their help in conducting the radiation experiments and Ms. Mehreen Saigal and Ms. Thejaswi B S for their overall help during the conduct of experiments.

## Authors’ contributions

VBS maintained the animals, performed and analyzed the experiments, and wrote the manuscript. KS and SN performed the irradiation procedure, DRN processed the samples for 16S rRNA sequencing and performed the downstream processing, and VBS and AJ analyzed the 16S rRNA sequencing data. KDM, TSM, KS, and SN provided significant comments and revisions to the article.

